# Benchmarking Phenotypic Clustering Algorithms via Empirically Calibrated Simulations: A Diagnostic Framework to Improve Biodiversity Assessment in Neglected Crops

**DOI:** 10.1101/2025.07.16.665063

**Authors:** Abdel Kader Naino Jika

**Affiliations:** University Abdou Moumouni, Faculty of Agronomy, Department of crops productions, Niamey, Niger

**Keywords:** Phenotypic Clustering, Simulation Benchmark, Biodiversity Assessment, Neglected and Underutilized Species (NUS), *Digitaria exilis*, Germplasm Management, PCA/ UMAP

## Abstract

Clustering algorithms are widely used for phenotypic characterization and germplasm management, particularly in neglected and underutilized species (NUS) that lack genomic resources. However, their performance under biologically realistic conditions remains poorly understood. Standard clustering methods commonly applied in crop research often assume distinct, isotropic, and homogeneous clusters—assumptions rarely satisfied in real-world NUS datasets.

We developed a biologically informed simulation framework, empirically calibrated with phenotypic data from West African fonio (Digitaria exilis), to benchmark the performance of eleven clustering algorithms under both idealized and realistic scenarios. Our simulations integrated heterogeneous trait distributions (normal, gamma), strong inter-trait correlations (up to r = –0.84), heteroscedasticity, and moderate population structure (Pst ≈ 0.15), as observed in fonio landraces. Each scenario was replicated 100 times, with clustering accuracy evaluated using Adjusted Rand Index (ARI), Normalized Mutual Information (NMI), Silhouette coefficient, and Davies–Bouldin index.

The results revealed consistently poor algorithm performance under realistic conditions (e.g., ARI < 0.07), including for widely used methods in NUS research such as K-means, GMM, and PAM. Performance markedly improved under idealized conditions, validating our simulation framework.

These findings highlight the risk of overinterpreting clustering outputs from weakly structured phenotypic datasets and expose key limitations in current biodiversity analysis practices—particularly those guiding plant genetic resource conservation programs. We provide an open-source R-based diagnostic tool, available on Zenodo 5(https://doi.org/10.5281/zenodo.15877863), to assist practitioners in selecting robust clustering approaches for germplasm management and pre-breeding in data-scarce crops.

## Introduction

Neglected and underutilized species (NUS) constitute a critical reservoir of agricultural biodiversity, essential for global food security, sustainable agriculture, and adaptation to climate change and ecological instability (Mabhaudhi et al., 2019; Padulosi et al., 2021). Despite their resilience and adaptive potential, many NUS—such as fonio (*Digitaria exilis*), a cereal cultivated in ecologically marginal environments—remain underrepresented in genomic databases due to limited funding and molecular resources (Bio et al., 2020; Vodouhè et al., 2022). Consequently, scientists and breeders often rely on phenotypic data as a primary resource for exploring genetic structure, managing germplasm collections, and informing breeding strategies (Pilling et al., 2020).

Yet, phenotypic analyses face fundamental challenges. These include subtle population differentiation and complex data distributions—characteristics often overlooked in methodological practice (Jombart et al., 2010; Odong et al., 2011). Fonio epitomizes this issue: historically vital to marginalized West African communities, it thrives in agroecological niches unsuitable for major cereals (Adoukonou-Sagbadja et al., 2007; Bio et al., 2020). In the absence of genomic profiling, phenotypic clustering is often the only available framework guiding biodiversity assessments and conservation interventions (Sogbohossou & Achigan-Dako, 2014).

Crucially, the interpretability of phenotypic clustering depends on assumptions that are rarely met in NUS datasets—namely, that clusters are discrete, isotropic, and internally homogeneous (Bouveyron et al., 2007; Jain et al., 1999). In practice, phenotypic traits in NUS often exhibit overlapping distributions, strong inter-trait correlations, heteroscedasticity, and low genetic differentiation (Pst ≈ 0.1–0.15), reflecting continuous gene flow and mixed mating systems (Vinh et al., 2010; Romano et al., 2016; Mahmoudi et al., 2024).

Despite these biological complexities, most benchmarking studies assume idealized conditions. This disconnect raises concerns about the validity of widely used clustering metrics—such as Adjusted Rand Index (ARI) and Normalized Mutual Information (NMI)—which are sensitive to violations of assumptions like balanced sample sizes, distinct separation, and homogeneous variance (Hubert & Arabie, 1985; Vinh et al., 2010). While methodological critiques have emerged (Romano et al., 2016), few empirical studies have systematically quantified how these biases affect clustering outcomes in realistic biological scenarios. This gap is structural, not merely technical.

Several popular algorithms illustrate this misalignment:

- **K-means** and **PAM** assume spherical clusters with equal variances—conditions rarely met in empirical datasets (Jain et al., 1999);
- **Ward’s method** favors compact, globular clusters but collapses under trait-driven asymmetries (Murtagh & Legendre, 2014);
- **Gaussian Mixture Models (GMM)** offer flexibility but still assume normality, often violated by real traits (McLachlan & Peel, 2000);
- **DBSCAN** accommodates arbitrarily shaped clusters but fails when densities vary across groups (Ester et al., 1996).

These algorithmic limitations have real-world consequences. Misclassification can result in the exclusion of locally adapted landraces or the loss of drought-resilient accessions—missed opportunities that are particularly costly in resource-limited settings. This structural misalignment directly undermines biodiversity conservation targets. As demonstrated by Arneth et al. (2020), climate pressures threaten to invalidate preservation strategies for NUS based on artificial phenotypic clusters—even when controlling for non-climatic pressures like habitat exploitation and ecological fragmentation.

To address this methodological gap, we introduce a biologically grounded simulation framework tailored to the statistical and ecological realities of neglected and underutilized species (NUS). Calibrated with empirical phenotypic data from West African fonio (Digitaria exilis), the framework enables rigorous benchmarking of eleven clustering algorithms— including K-means, PAM, Ward’s method, GMM, DBSCAN, HDBSCAN, Spectral clustering, Fuzzy C-means, Affinity Propagation, Self-Organizing Maps (SOM), and Two-Step clustering—under both idealized and realistic conditions. The simulated datasets reproduce key real-world complexities: heterogeneous trait distributions (normal and gamma), strong inter-trait correlations (up to r = –0.84), heteroscedastic variances, and moderate population structure (Pst ≈ 0.15). We evaluate algorithm performance across 100 simulation replicates using ARI, NMI, Silhouette coefficient, and Davies–Bouldin index, supplemented by dimensionality reduction tools such as PCA and UMAP to assess the geometric structure of clusters.

Our central hypothesis is that apparent clusters in phenotypic datasets of NUS with modest genetic structure often arise as statistical artefacts driven by methodological constraints, rather than reflecting genuine biological partitions. By testing this systematically, our study offers a reproducible, R-based diagnostic tool to evaluate clustering validity before drawing evolutionary or agronomic conclusions.

We advocate for a paradigm shift: empirically calibrated simulations should replace idealized Gaussian models as the standard for evaluating clustering performance in NUS research. This shift prioritizes biological realism over computational convenience—particularly in regions such as sub-Saharan Africa, where genotyping is often inaccessible, while phenotyping remains more feasible and cost-effective. The tool is open-source, modular, and designed for reproducibility and cross-species adaptability. To maximize generalizability, its parameters were not only calibrated from fonio, but also designed to accommodate empirical data from other NUS available in the literature.

This article thus serves as the methodological cornerstone of a broader research agenda aimed at systematically benchmarking clustering approaches for biodiversity assessment across diverse underutilized taxa. It promotes a phenotype-first paradigm in agrobiodiversity research—one that centers ecological realism, supports data-driven decision-making, and advances conservation and breeding efforts in resource-limited settings.

## Materials and Methods

### Empirical Data and Trait Architecture

This study draws upon agromorphological trait data from 180 *Digitaria exilis* (fonio) landraces characterized by Bio et al. (2020), spanning three geographically distinct groups (n = 91, 43, and 46). Eight traits were selected for their agronomic relevance and quantitative tractability: plant height (PHT), number of internodes (NIN), flowering time (FLO), maturity time (MAT), number of grains per inflorescence (NGR), grain yield per plant (GRY), harvest index (HI), and thousand-seed weight (TSW). Trait distributions were predominantly normal, with the exception of GRY, which followed a gamma distribution. Empirical trait correlations (range: *r* = –0.84 to 0.77) were encoded in a near-positive definite correlation matrix (Higham, 2002), and group-specific variances were retained to reproduce heteroscedastic structures in simulations.

### Simulation Framework

Two simulation scenarios were implemented to assess clustering robustness:

1. Realistic Scenario: Phenotypic datasets (n = 180) were synthetically generated using a multivariate transformation pipeline:

- **Correlated base generation:** Traits simulated with MASS::mvrnorm() to match empirical covariance structure.
- **Distributional alignment:** Gamma-distributed traits (e.g., GRY) transformed from normal deviates via quantile matching.
- **Population structure:** Group-level covariance matrices were iteratively tuned to achieve a target phenotypic differentiation of Pst ≈ 0.15 (mean realized: 0.16 ± 0.001).
- **Heteroscedasticity:** Trait-wise standard deviations were scaled per group using empirical multipliers (e.g., Pop1 GRY SD × 0.8).
2. Idealized Control: A benchmark dataset featuring:

- Balanced group sizes (n = 50 per group),
- High between-group differentiation (Pst = 0.80),
- Independent traits (identity matrix),
- Homoscedastic variances.

### Clustering Algorithms

We benchmarked eleven clustering methods representing major analytical paradigms:

- **Partitioning:** K-means, PAM;
- **Hierarchical:** Ward’s method;
- **Model-based:** Gaussian Mixture Models (GMM);
- **Density-based:** DBSCAN, HDBSCAN;
- **Graph-theoretic:** Spectral clustering;
- **Affinity-based:** Affinity Propagation;
- **Fuzzy clustering:** Fuzzy C-means;
- **Neural-network-based:** Self-Organizing Maps (SOM);
- **Hybrid scalable:** Two-Step clustering.

For parametric methods, the number of clusters (*k* = 3) was fixed based on known population structure (Bio et al., 2020). Non-parametric methods relied on data-driven heuristics. Parameter tuning followed best-practice guidelines (e.g., DBSCAN’s eps set to 90th percentile of kNN distances; minPts = 5).

### Evaluation metrics

Clustering outcomes were evaluated using four complementary metrics:

- **Adjusted Rand Index (ARI):** Measures agreement between true and predicted labels, corrected for chance (Hubert & Arabie, 1985).
- **Normalized Mutual Information (NMI):** Quantifies shared information independent of label permutations (Vinh et al., 2010).
- **Silhouette Coefficient:** Captures internal cohesion and external separation (Rousseeuw, 1987).
- **Davies–Bouldin Index:** Evaluates cluster compactness relative to separation (Davies & Bouldin, 1979).

A dispersion index (betadisper; Anderson, 2006) was also computed to assess within true population variability relative to group centroids.

### Statistical analysis and reproducibility

Each scenario was replicated 100 times. Performance metrics were summarized using means and standard deviations; robustness was quantified as the inverse of inter-replicate variance. Pairwise comparisons employed Dunn’s tests with FDR correction (α = 0.01). Visualization of clustering validity employed Principal Component Analysis (PCA) and Uniform Manifold Approximation and Projection (UMAP) (Jolliffe & Cadima, 2016; McInnes et al., 2018).

All R scripts (version 4.3.0), including simulations, clustering pipelines, and validation routines, are publicly archived on Zenodo (DOI: https://doi.org/10.5281/zenodo.15877863). Reproducibility is ensured through modular scripting:

- simulate_populations.R: Synthetic data generation
- clustering_benchmark.R: Algorithm implementations
- validation_tools.R: Metrics and dispersion
- result_analysis.R: Statistical comparison and visualization

## Results

### Clustering Performance Under Realistic Phenotypic Conditions

Under phenotypic conditions empirically calibrated from West African fonio landraces (Pst ≈ 0.16), all eleven tested clustering algorithms exhibited uniformly low performance across validation metrics (Table 1, Figure 1). The Adjusted Rand Index (ARI) ranged from –0.006 (HDBSCAN) to 0.053 (GMM), indicating near-random agreement with true groupings. Normalized Mutual Information (NMI) values were similarly low (maximum: 0.083, Affinity Propagation), suggesting poor information retention. Silhouette coefficients exceeded the interpretability threshold (0.25) in only one case (DBSCAN: 0.379), while Davies–Bouldin indices consistently exceeded 1.2, signifying poor inter-cluster separation.

**Figure 1.**
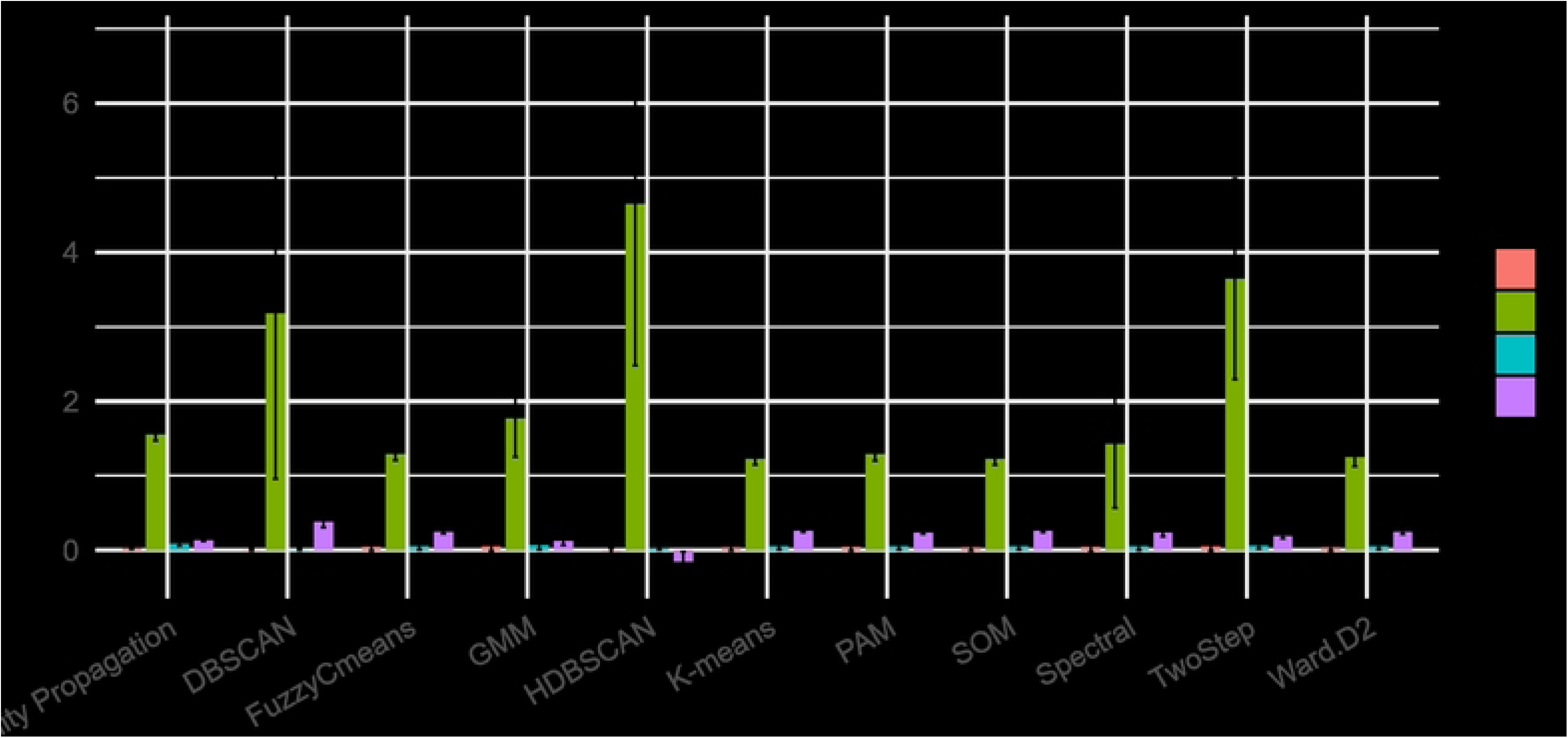
Mean performance (± SD) of eleven clustering methods across ARI, NMI, Silhouette, and Davies–Bouldin indices in simulations calibrated with empirical fonio trait data (Pst ≈ 0.16).

**Table 1.** Mean Clustering Metrics Under Realistic Conditions (± SD; n = 100 unless noted)

| Méthode | ARI | NMI | Silhouette | Davies-Bouldin |
| --- | --- | --- | --- | --- |
| GMM | 0.053 ± 0.064 | 0.070 ± 0.062 | 0.127 ± 0.059 | 1.768 ± 0.516 |
| TwoStep | 0.051 ± 0.058 | 0.063 ± 0.048 | 0.188 ± 0.032 | 3.641 ± 1.347 |
| Spectral | 0.045 ± 0.066 | 0.055 ± 0.052 | 0.240 ± 0.057 | 1.429 ± 0.862 |
| PAM | 0.045 ± 0.050 | 0.058 ± 0.050 | 0.241 ± 0.025 | 1.289 ± 0.090 |
| Fuzzy C-means | 0.042 ± 0.047 | 0.055 ± 0.048 | 0.244 ± 0.023 | 1.288 ± 0.083 |
| SOM | 0.042 ± 0.045 | 0.057 ± 0.048 | 0.262 ± 0.022 | 1.223 ± 0.079 |
| K-means | 0.041 ± 0.044 | 0.056 ± 0.047 | 0.263 ± 0.021 | 1.223 ± 0.077 |
| Ward.D2 | 0.040 ± 0.044 | 0.056 ± 0.045 | 0.243 ± 0.029 | 1.249 ± 0.124 |
| Affinity<br>Propagation | **0.031 ± 0.019** | **0.083 ± 0.028** | 0.132 ± 0.011 | 1.556 ± 0.087 |
| DBSCAN | 0.007 ± 0.011 | 0.009 ± 0.008 | **0.379 ± 0.073** | 3.184 ± 2.226 |
| HDBSCAN<br>(n=89) | -0.006 ± 0.031 | 0.033 ± 0.026 | -0.153 ± 0.132 | **4.648 ± 2.169** |

Importantly, these results were consistent across 100 simulation replicates per algorithm (except HDBSCAN: n = 89 due to convergence issues), reinforcing the reliability of observed patterns. Despite modest differences between methods, no algorithm achieved a biologically meaningful clustering outcome under realistic trait complexity and moderate population differentiation.

### Performance Under Idealized Conditions

In the idealized scenario featuring high differentiation (Pst ≈ 0.80), independent traits, equal sample sizes, and homoscedasticity, algorithm performance dramatically improved (Table 2, Figure 2). Parametric methods such as GMM, Fuzzy C-means, and K-means achieved near-perfect ARI (>0.97) and NMI (>0.96) values, confirming both the integrity of the simulation pipeline and the sensitivity of these algorithms to favorable statistical conditions. Silhouette coefficients (∼0.39) and Davies–Bouldin indices (∼1.05) indicated well-separated, coherent clusters.

**Figure 2.**
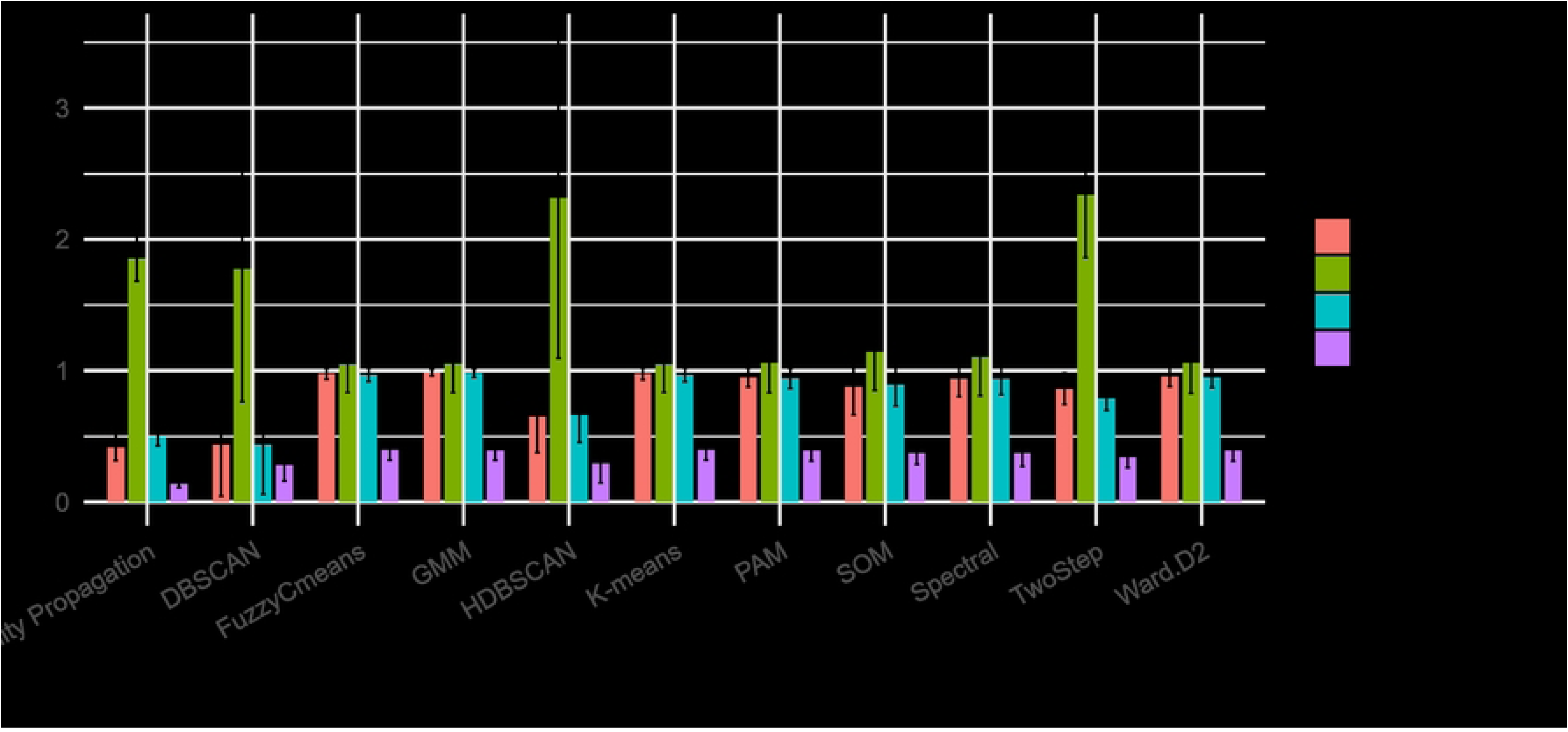
Validation of simulation pipeline and algorithmic capacity under favorable statistical assumptions: independent traits, strong structure, and homoscedasticity.

**Table 2.** Mean Clustering Metrics Under Ideal Conditions (± SD; n = 100)

| Méthode | ARI | NMI | Silhouette | Davies-Bouldin |
| --- | --- | --- | --- | --- |
| **GMM** | 0.986 ± 0.026 | 0.981 ± 0.033 | 0.394 ± 0.077 | 1.051 ± 0.219 |
| **Fuzzy C-means** | 0.976 ± 0.042 | 0.968 ± 0.052 | 0.394 ± 0.076 | 1.048 ± 0.215 |
| **K-means** | 0.975 ± 0.045 | 0.968 ± 0.053 | 0.394 ± 0.076 | 1.047 ± 0.213 |
| **Ward.D2** | 0.955 ± 0.076 | 0.949 ± 0.077 | 0.390 ± 0.080 | 1.061 ± 0.234 |
| **PAM** | 0.948 ± 0.073 | 0.941 ± 0.077 | 0.390 ± 0.080 | 1.058 ± 0.227 |
| **Spectral** | 0.937 ± 0.134 | 0.936 ± 0.119 | 0.372 ± 0.101 | 1.102 ± 0.293 |
| **SOM** | 0.875 ± 0.212 | 0.894 ± 0.164 | 0.371 ± 0.086 | 1.142 ± 0.292 |
| **TwoStep** | 0.862 ± 0.119 | 0.790 ± 0.093 | 0.340 ± 0.079 | 2.339 ± 0.478 |
| **HDBSCAN** | 0.652 ± 0.276 | 0.663 ± 0.210 | 0.292 ± 0.147 | 2.317 ± 1.223 |
| **DBSCAN** | 0.436 ± 0.391 | 0.437 ± 0.378 | 0.282 ± 0.122 | 1.775 ± 1.011 |
| **Affinity Propagation** | 0.415 ± 0.101 | 0.496 ± 0.069 | 0.136 ± 0.027 | 1.856 ± 0.174 |

By contrast, non-parametric methods (e.g., DBSCAN, HDBSCAN, Affinity Propagation) showed lower mean performance and higher variance, suggesting reduced reliability even under ideal conditions. DBSCAN failed to converge in 4% of replicates due to poor density parameter fits.

### Visual diagnostics reveal latent structure

While numerical metrics consistently suggested low clustering validity under realistic scenarios, dimension reduction techniques (PCA and UMAP) revealed subtle groupings in certain algorithms (figure 3, S1 to S11). Notably, DBSCAN and Affinity Propagation identified structure congruent with known groupings, despite low ARI/NMI scores. PCA explained >80% of variance on the first two axes, underscoring biologically relevant structure that escapes conventional metrics. This discordance highlights the limitations of relying solely on statistical indices when interpreting weakly differentiated datasets.

**Figure 3.**
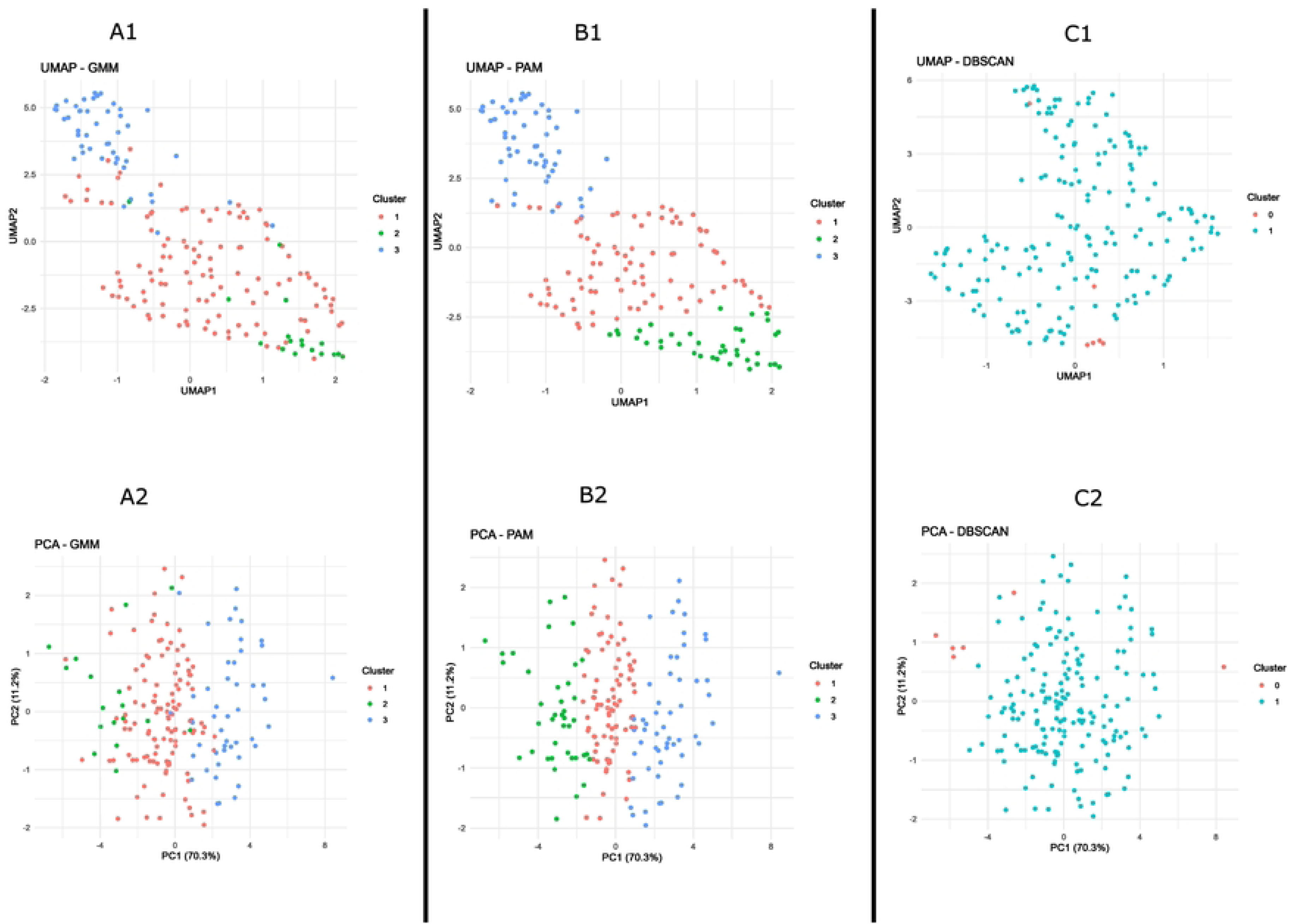
PCA and UMAP Reveal Latent Structure Beyond Numerical Metrics *Visual representation of clustering patterns across dimensionality reduction methods, indicating underlying structure undetected by ARI and NMI*.

## Discussion

Our study presents a rigorous and biologically informed evaluation of clustering methods applied to phenotypic datasets from neglected and underutilized species (NUS), exemplified by fonio. It reveals several critical insights:

1. **Fundamental Methodological Mismatch:** Under realistic phenotypic simulations (Pst ≈ 0.16; group sizes 91, 43, 46), all tested clustering algorithms—ranging from partition-based (K-means, PAM) to model-based (GMM) and density-based methods (DBSCAN, HDBSCAN)— yielded uniformly poor agreement with true group labels (ARI < 0.06; NMI < 0.09). This quantitatively demonstrates the fundamental limitations noted by Jombart et al. (2010) and Odong et al. (2011), extending their qualitative insights to simulation-based rigor. The reliance on assumptions of discrete, isotropic, homogeneous clusters in these algorithms is misaligned with the overlapping, correlated, heteroscedastic, and weakly differentiated structures typical of NUS phenotypic data (Bio et al., 2020; Mahmoudi et al., 2024).
2. **ARI/NMI Fail to Detect Subtle Biological Structure:** Despite consistently low ARI and NMI scores, PCA and UMAP visualizations revealed coherent group separation in some clustering outputs—capturing over 80% of variance on the first two axes. This uncoupling reinforces critiques by Romano et al. (2016) and Vinh et al. (2010), showing that conventional metrics inherently underestimate biologically meaningful differentiation when clusters are continuous or overlapping. We argue these low scores should be viewed as signals of underlying biological complexity, rather than algorithmic failure.
3. **Visualization Alone Is Insufficient — Diagnostics Must Combine Metrics and Plots** Our findings underscore the necessity of integrating qualitative (PCA/UMAP) and quantitative diagnostics. Despite pronounced visual structure in DBSCAN and Affinity Propagation outputs, reliance solely on ARI/NMI would reject these methods. This aligns with critiques in single-cell genomics and suggests biodiversity analyses require multidimensional validation protocols (McInnes et al., 2018; Jolliffe & Cadima, 2016).
4. **Trade-Off Between Stability and Accuracy:** Analyzing robustness (inverse ARI variance) exposes a critical practical trade-off: high-performing but unstable methods (e.g., GMM) versus more stable, interpretable algorithms (e.g., PAM, Ward’s hierarchical). This finding supports prioritizing reproducibility and interpretability in applied NUS research—a stance aligned with Pilling et al. (2020) and Vodouhè et al. (2022).
5. **Toward Data-Driven Clustering via Empirical Simulation:** Crucially, our framework demonstrates the value of context-specific simulation. By parameterizing distributions (normal, gamma, binomial), sample imbalance, correlation structures, heteroscedasticity, and group number, researchers can identify clustering methods most robust to specific dataset properties and assumption violations. This moves phenotypic clustering from a one-size-fits-all approach to a tailored, data-driven strategy.
6. **Need for Novel Clustering Validation Metrics:** The observed mismatch between conventional validation metrics and visual/biological structure highlights the urgent necessity for new indices—designed for weakly structured NUS data. Such metrics must integrate geometric separation, internal cohesion, and be robust to non-normality, unbalanced designs, heteroscedasticity, and nonlinear correlations.
7. **Mapping Performance Across Population Differentiation Gradient:** Future work should focus on how clustering detectability varies with phenotypic differentiation. As Jombart et al. (2010) suggest, there might exist Pst thresholds below which conventional clustering is unreliable. Empirical mapping across Pst gradients would enable researchers to distinguish diffuse biological structure from true method failure and develop general interpretative frameworks.
8. **Broad Impact and Future Extensions:** Our study challenges current NUS phenotypic research paradigms—routinely reliant on K-means, GMM, or DBSCAN. Instead, we propose a robust methodological reform grounded in empirically calibrated simulations, combined validation diagnostics, and context-aware algorithm selection. This paradigm is generalizable to NUS with varying sample sizes, trait distributions, complex correlations, and population structures. Future extensions should also incorporate missing data mechanisms (MCAR/MAR), extreme sample imbalance, continuous differentiation gradients, and integration with genomic validation (Elshire et al., 2011).

## Conclusion

This study demonstrates that conventional clustering algorithms systematically underperform when applied to realistically simulated phenotypic data from neglected and underutilized species (NUS), such as fonio. These results do not merely reflect methodological shortcomings, but expose a fundamental misalignment between algorithmic assumptions and the biological realities of NUS datasets—characterized by overlapping traits, heteroscedasticity, and modest differentiation.

Critically, we show that low ARI and NMI scores can obscure meaningful structure, which is nevertheless revealed by visual diagnostics such as PCA and UMAP. This highlights the need to treat clustering as an exploratory rather than confirmatory tool, and to complement numeric metrics with qualitative evaluation.

Our biologically calibrated simulation framework offers a reproducible, flexible diagnostic platform to benchmark clustering performance under realistic conditions. It enables researchers to align methodological choices with empirical data properties, supporting more robust and interpretable phenotypic stratification.

Looking ahead, future research should develop alternative validation metrics better suited to continuous and weakly structured biological data, and investigate how clustering performance varies across the full gradient of population differentiation. Integrating missing data patterns and genomic validation will further refine this approach, strengthening its utility for conservation, breeding, and germplasm management in data-scarce crops.

## Supporting Informations

- S1 Figure. Affinity propagation clustering results under realistic conditions (Pst ≈ 0.15) using PCA (a) and UMAP (b)
- S2 Figure. Fuzzy c means clustering results under realistic conditions (Pst ≈ 0.15) using PCA (a) and UMAP (b)
- S3 Figure. HDBSCAN clustering results under realistic conditions (Pst ≈ 0.15) using PCA (a) and UMAP (b)
- S4 Figure. K-means clustering results under realistic conditions (Pst ≈ 0.15) using PCA (a) and UMAP (b)
- S5 Figure. SOM clustering results under realistic conditions (Pst ≈ 0.15) using PCA (a) and UMAP (b)
- S6 Figure. Spectral clustering results under realistic conditions (Pst ≈ 0.15) using PCA (a) and UMAP (b)
- S7 Figure. TwoStep clustering results under realistic conditions (Pst ≈ 0.15) using PCA (a) and UMAP (b)
- S8 Figure. Ward clustering results under realistic conditions (Pst ≈ 0.15) using PCA (a) and UMAP (b)
- S9 Figure. Affinity propagation clustering results under ideal conditions using PCA (a) and UMAP (b)
- S10 Figure. DBSCAN clustering results under ideal conditions using PCA (a) and UMAP (b)
- S11 Figure. Fuzzy C Means clustering results under ideal conditions using PCA (a) and UMAP (b)

